# Multi-Site Statistical Mapping of Along-Tract Microstructural Abnormalities in Bipolar Disorder with Diffusion MRI Tractometry

**DOI:** 10.1101/2023.08.17.553762

**Authors:** Leila Nabulsi, Bramsh Q. Chandio, Genevieve McPhilemy, Fiona M. Martyn, Gloria Roberts, Brian Hallahan, Udo Dannlowski, Tilo Kircher, Benno Haarman, Philip Mitchell, Colm McDonald, Dara M. Cannon, Ole A. Andreassen, Christopher R. K. Ching, Paul M. Thompson

## Abstract

Investigating alterations in brain circuitry associated with bipolar disorder (BD) may offer a valuable approach to discover brain biomarkers for genetic and interventional studies of the disorder and related mental illnesses. Some diffusion MRI studies report evidence of microstructural abnormalities in white matter regions of interest, but we lack a fine-scale spatial mapping of brain microstructural differences along tracts in BD. We also lack large-scale studies that integrate tractometry data from multiple sites, as larger datasets can greatly enhance power to detect subtle effects and assess whether effects replicate across larger international datasets. In this multisite diffusion MRI study, we used BUndle ANalytics (BUAN, Chandio 2020), a recently developed analytic approach for tractography, to extract, map, and visualize profiles of microstructural abnormalities on 3D models of fiber tracts in 148 participants with BD and 259 healthy controls from 6 independent scan sites. Modeling site differences as random effects, we investigated along-tract white matter (WM) microstructural differences between diagnostic groups. QQ plots showed that group differences were gradually enhanced as more sites were added. Using the BUAN pipeline, BD was associated with lower mean fractional anisotropy (FA) in fronto-limbic, interhemispheric, and posterior pathways; higher FA was also noted in posterior bundles, relative to controls. By integrating tractography and anatomical information, BUAN effectively captures unique effects along white matter (WM) tracts, providing valuable insights into anatomical variations that may assist in the classification of diseases.

## I. Introduction

Mental health disorders affect more than 1 billion people worldwide [1]. Psychiatric conditions contribute to 7% of the global burden of disease, measured using disability-adjusted life years, and account for 19% of all years lived with disability [1]. In the United States alone, serious mental illnesses, including bipolar disorder (BD), directly affect nearly one in 25 adults [1]. It is crucial to reliably characterize the neural substrates involved in mental health disorders. Such characterization is essential to accelerate the discovery of brain biomarkers for genetic and interventional studies, for accurate targeting of treatments, outcome prediction, and modeling inter- and intra-individual disease variability [1]. Among the affective disorders, BD has increasingly been recognized as a *dysconnection syndrome* with deficits in both gray and white matter components of brain circuits involved in regulation of emotion and reward processing [2]. Contemporary neurobiological theories propose that BD is associated with alterations in fronto-limbic brain regions, their connections, as well as impaired interactions with brain areas traditionally implicated in anxiety, fear, and cognitive control [2]. These theories suggest that dysfunction in these fronto-limbic circuits underlies the emotional and cognitive dysregulation characteristic of the disorder [2]. Current models of BD emphasize the importance of understanding the interplay between gray and white matter components involved in emotion regulation. Prior research using structural brain MRI, such as voxel-based morphometry (VBM) [3], has attempted to characterize white matter differences in BD [4], but T1-weighted imaging offers limited contrast within the white matter. Diffusion MRI (dMRI) offers advanced metrics of white matter microstructure, but these approaches have not yet provided a detailed spatial mapping of microstructural differences in BD [5]. Instead, most dMRI studies of BD have used techniques that average measures along tracts, for example region-of-interest (ROI) analyses, such as tract-based spatial statistics (TBSS), and more recently, network-based approaches [6]. Identifying abnormalities in specific segments of white matter bundles in brain disorders is crucial. Precisely localizing damage on a tract can aid in accurate diagnosis and differential diagnosis of various brain disorders. This information is invaluable for tracking disease progression over time and tailoring treatment plans to the specific affected areas. Understanding the functional consequences of tract damage in these sections helps in designing targeted intervention strategies. Additionally, this knowledge may support personalized medicine and provide prognostic value, informing patients and healthcare providers about potential outcomes.

The literature on white matter connectivity in BD reveals deviations in white matter microstructure in key association pathways, predominantly characterized by limbic dysconnectivity, which serves as a distinctive hallmark of the illness [3, 7, 8, 9, 10, 11]. These white matter projections consist of tracts that convey signals to and from cortical regions responsible for regulating the emotional, cognitive, and behavioral dimensions of BD [3, 7, 8, 9, 10, 11]. Research on white matter structure in BD reveals that abnormalities likely extend beyond the anterior fronto-limbic pathways, encompassing a broader spectrum of connections. Notably, alterations have been reported in superior longitudinal (arcuate) and inferior *fasciculi*, inferior fronto-occipital gyrus, posterior thalamic radiation, and the internal capsule in BD. Furthermore, compromised white matter connectivity has been documented within posterior white matter tracts, such as the superior occipital white matter and cerebellum [3, 7, 8, 9, 10, 11], which may act as compensatory mechanisms in response to disrupted frontal connectivity. These changes collectively contribute to cognitive impairments and emotional dysregulation in individuals with BD. White matter deficits might arise from a lack of coherence, variations in the quantity or density of white matter tracts, or potential disruptions in myelin or axonal integrity. White matter bundle segmentation has greatly advanced recently, due to the progress in tractography techniques based on diffusion MRI. These advancements include automated bundle segmentation in brain research, enabling the study of neural pathways yielding valuable insights into the microstructural properties of white matter that differ between diseased and healthy brains. Here, we used BUndle ANalytics (BUAN) [12, 3], a recently developed approach for tractography analysis, to reconstruct refined anatomic maps along the length of WM bundles extracted from whole-brain tractograms in BD and healthy controls. BUAN is a step forward from commonly applied techniques that average metrics along the full length of the tract. BUAN has recently been applied to visualize the profile of microstructural abnormalities on 3D models of fiber tracts, yielding fine-scale maps of the effects of Parkinson’s disease [12], mild cognitive impairment (in ADNI3; [13]), and aging [14]. BUAN is also freely available in DIPY - https://dipy.org/documentation/1.5.0/interfaces/buan_flow/. In a pilot study, using the BUAN pipeline, BD (N=38) was associated with lower mean fractional anisotropy (FA) in fronto-limbic and interhemispheric pathways and higher mean FA in posterior bundles relative to healthy controls (N=49) [15]. Small to moderate effect sizes are the norm for BD and many other related mental illnesses. Although informative, a larger-scale multi-center, big-data approach would maximize statistical power, which is crucial for finding robust, reliable effects on the brain’s white matter connectivity and determining how generalizable effects are across independent cohorts. In this study, we combined data from 6 independent scan sites to maximize statistical power and assess evidence for previously reported features of microstructural deficits in BD, with the anatomically improved precision conferred by BUAN, relative to controls. We hypothesized that people with BD would exhibit the greatest alterations in limbic, interhemispheric, and cerebellar white matter projections.

## II. Methods

We analyzed cross-sectional 3D diffusion-weighted MRI data from 6 independent scanning centers (average age 36; range 18-64 years), from 148 individuals with BD (age: 36.8±13.5 y; 48% female) and 259 psychiatrically healthy controls (age: 37.7±13.7 y; 57% female). Each site had unique acquisition parameters (gradient directions range=30-64, and *b*-values=1000-1200 s/mm^2^). For details of acquisition parameters and socio-demographic data from these sites, please see [16-20]. All raw data was visually inspected to check for problems in image acquisition, including movement, susceptibility, and noise artifacts. Signal-to-noise ratio was checked for all scans to ensure quality of diffusion signal and suitability for tractography. A total of 22 subjects were excluded from our analysis due to poor quality scans. The remaining scans were corrected for subject motion, including rotating the *b*-matrix, and corrected for eddy-current distortions (ExploreDTI v4.8.6) [21]. To account for crossing fibers within voxels, we used a deterministic (non-tensor) constrained spherical deconvolution (CSD) algorithm [22] (ExploreDTI). We used CSD, a more anatomically meaningful approach than standard tensor-based algorithms, that better characterizes the underlying microstructural organization of the fiber bundles. Each site preprocessed their diffusion data and the whole-brain tractogram was qualitatively and quantitatively examined to guarantee successful reconstruction of streamlines for each subject. Diffusion eigenvector estimation was performed using the robust estimation of tensor by outlier rejection (RESTORE) approach [23]. For all sites, fiber tracking commenced in each voxel, continued with 1-mm step size, 2 × 2 x 2 mm seed point resolution, >30° angle curvature threshold, and 20–300 mm length, and terminated at a minimum fractional anisotropy (FA) value of 0.2. FA, a measure of microstructural tissue organization, was calculated at each voxel (in this proof-of-concept study, we focus on FA due to space constraints, but the methods are applicable to other tensor and multicompartmental models computed at each point along tracts). Each subject’s FA map was affinely registered to an FA template in MNI space using the registration framework provided in DIPY [24] with an implementation similar to ANTs [25]. Individual whole-brain tractograms underwent streamline-based linear registration (SLR) [26] to a bundle atlas template in MNI space [27]. 37 bundles were extracted from each subject’s whole-brain tractogram using the auto-calibrated version of RecoBundles and a standard WM tract atlas [28]. Next, we employed the BUAN tractometry pipeline to identify significant microstructural group differences at specific locations along the trajectory of each tract. A tract profile was generated for each extracted bundle, where fractional anisotropy (FA) values were assigned to all points of the streamlines within the bundle for each subject (*Fig. 1*). BUAN differs from many traditional tractometry methods, which simplify each tract to a single mean streamline [29], [30]. Instead, BUAN uses all points on all streamlines within the bundle to create the bundle profile [12, 13, 31]. To achieve this, each point on every streamline was assigned a segment number based on its proximity to the centroid of the model bundle (Euclidean distance), derived from an atlas. Each subject’s bundle profile consisted of 100 segments spanning the length of the tract. Data from the two groups were analyzed using linear mixed models (LMMs). For each bundle type, we fitted 100 linear mixed models, with one model per segment along the tract’s length. In these LMMs, fractional anisotropy (FA) was defined as the response variable, while group, age, and sex were considered fixed effect terms. To account for correlations among data points on the same streamline within a segment for a particular subject, as well as site-specific variations, subject and site were included as random effects terms. The use of random effects models ensured proper handling of these correlations. In this study, we identified specific segments within the bundles that exhibited differences in FA, a microstructural measure, between individuals with BD and the control group. To correct for multiple comparisons, we implemented False Discovery Rate (FDR) correction [32]. This correction was applied within each individual tract, considering the presence of 100 segments per tract. The FDR-adjusted threshold (\**pFDR*<0.05), in green in *Fig 1*., was used to account for multiple comparisons while considering the non-independence of segments within a tract. Additionally, we applied FDR across the entire brain, i.e., by pooling p-values across all 36 tracts, and the new FDR threshold (\**pFDR*=0.03) did not alter our main findings, shown in *Fig 2*.

**Figure 1.**
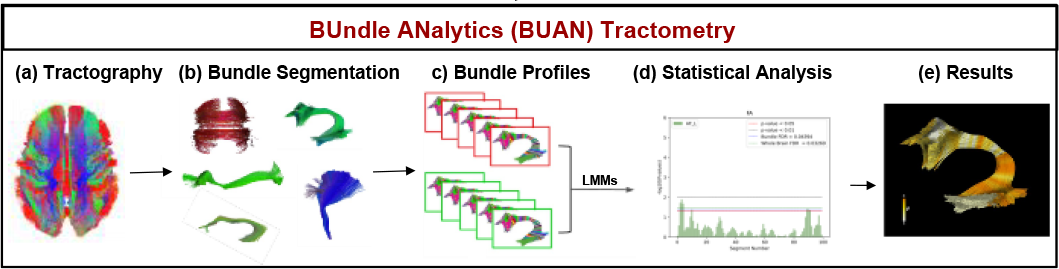
BUAN tractometry pipeline. (a) Whole-brain tractograms are reconstructed from brain dMRI scans using tractography; (b) Individual white matter tracts are extracted; (c) Bundle profiles are created from the sets of extracted bundles from patients and controls; here they are represented as 100 segments along the length of the tracts. (d) Bundle profiles are analyzed using LMMs, yielding a statistic of group difference in microstructural measures (such as FA) at each segment along the tracts. (d) These *p*-value profiles are displayed to highlight significant group differences from controls along the length of the tracts, which are FDR-corrected. (e) The *p*-values can be textured back onto the 3D models of the tracts.

**Figure 2.**
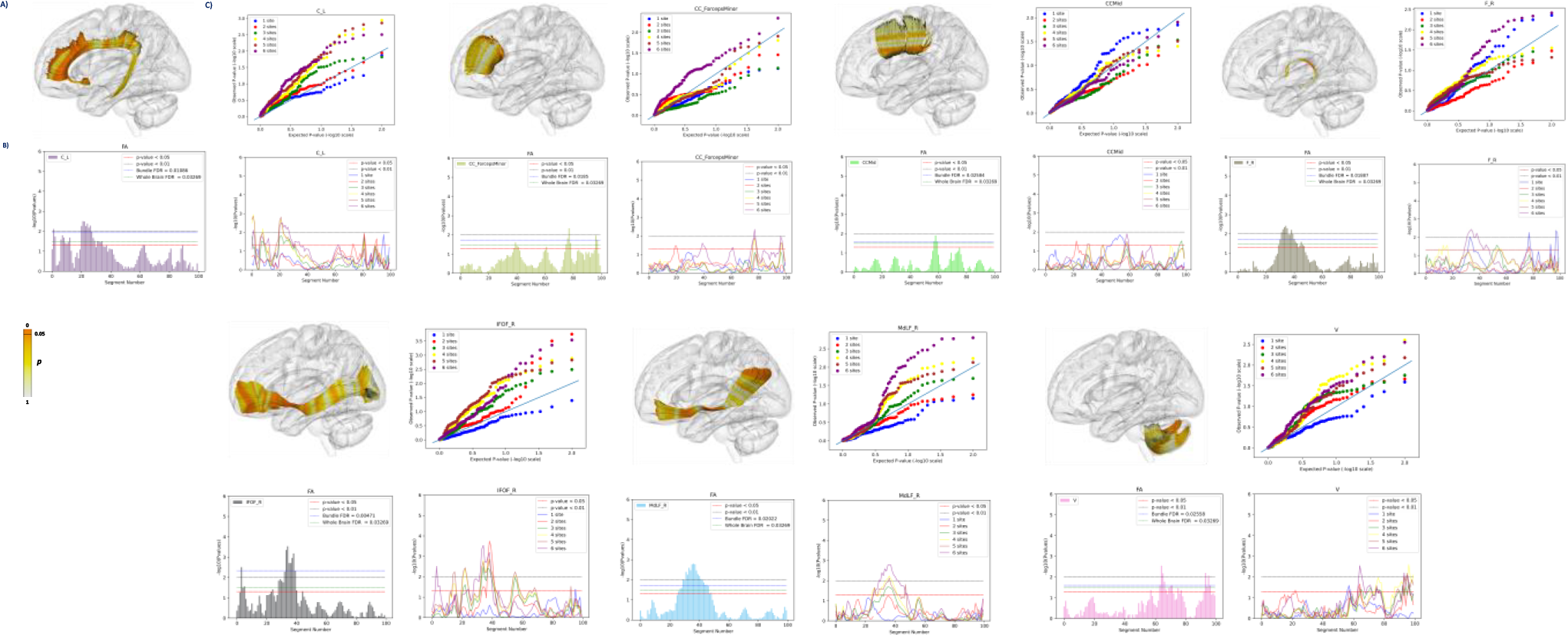
Along-tract microstructure (FA) alterations localized in BD. Compared to healthy controls, participants with BD exhibited altered FA in several bundles. Some are depicted here: C=cingulum; CC=corpus callosum; F=fornix; IFOF=inferior fronto-occipital fasciculus; MdLF=medial longitudinal fasciculus; V=vermis; L=left; R=right; Mid=middle; FA=fractional anisotropy. A) Significant *p*-values are shown in *dark orange* on each bundle. B) In each main plot, on the *x*-axis, segments along the length of the tract are shown and the *y*-axis shows the negative logarithm of the *p*-values. *P*-values between or above the two horizontal lines on the plot imply nominally significant (uncorrected) group differences at that location along the tract. The FDR adjusted threshold is plotted in blue; the whole-brain FDR threshold for each tract is plotted in green. The QQ plots in C) show that adding sites generally boosts effect size, for tracts where effects are identified.

## III. Results

Our findings suggest that microstructural abnormalities in BD may not be restricted to the white matter regions considered to support emotional regulation in the brain, such as fronto-limbic and inter-hemispheric connections. Instead, these abnormalities may also affect posterior projections. After multiple comparisons correction, compared to controls, the BD group exhibited lower FA within localized regions of the *cingulum* and the *fornix*. Additionally, regions within the *corpus callosum* (middle portion and *forceps minor*) and the fronto-parietal tract showed lower FA in the BD group. Several long-range intra-hemispheric white matter tracts exhibited lower FA in BD compared to controls in localized segments, and specifically the extreme capsule, the arcuate, uncinate, inferior-fronto occipital, medial- and middle-longitudinal *fasciculi*. Of note, the latter also exhibited higher FA in localized segments. Lower FA was detected in regions of the occipital-parietal tract and the vermis, and higher FA was seen in segments of the optic radiation, all relative to controls. Finally, in BD, white matter segments of the cortico-spinal tract, spino-thalamic and the medial lemniscus pathways exhibited lower FA compared to controls. For significant segments, please see *Fig. 2*.

## IV. Conclusions

In this multicenter study we applied the BUAN tractometry pipeline to map, extract, and visualize the effects of BD on the white matter tracts of the brain. We found significant microstructural differences in localized regions of fronto-limbic and inter-hemispheric pathways. Significant segments within posterior pathways and the brainstem also differed in the BD group compared to controls in terms of white matter microstructure. In BD, these effects were not evenly distributed across the length of the tract but appeared to be localized to specific segments, which may be related to underlying vulnerabilities in those specific regions of the tracts. Previous white matter studies in ENIGMA have reported overlapping effects across related mental illnesses [5]; BUAN may help to better quantify the overlapping or distinct points within and across disorders. Furthermore, adding more samples to this analysis improved effect size estimates for each tract. Abnormal FA in anterior portions within the cingulum is consistent with the regions where activations are detected in emotion-related fMRI studies in BD [33]. Morphological and network-based connectivity studies have reported deficits in FA in the limbic right anterior and subgenual cingulate cortex [34]. We found lower FA in the corpus callosum (middle portion and *forceps minor*) in BD. Alterations in the corpus callosum have been consistently found in structural, diffusion and functional MR studies of BD, particularly within the anterior horn, responsible for connecting bilateral prefrontal and limbic regions [34]. Morphological disturbances in the corpus callosum suggest inter-hemispheric structural dysconnectivity in BD. We also found alterations in segments within the fornix in BD. The fornix is interconnected with other brain regions involved in cognitive and emotional processing, such as the prefrontal cortex and the amygdala; whose abnormalities have been identified in prior ENIGMA BD studies [34]. Fornix abnormalities in BD may disrupt the flow of information between these regions, and impact cognitive and emotional functions, causing difficulties with attention, executive functions, and decision-making, as well as problems with emotion recognition and regulation [16]. Functional studies also highlight frontoparietal dysconnectivity in BD [35].

The frontoparietal network plays a crucial role in cognitive processes such as attention, working memory, and executive functions; disruptions in this network may contribute to cognitive impairments commonly observed in this disorder, including difficulties with attentional control, problem-solving, and decision-making. The frontoparietal network is also involved in emotion regulation and in integrating emotional and cognitive information. Dysconnectivity within this network may result in difficulties regulating emotions, leading to mood instability and emotional dysregulation frequently experienced by individuals with BD [35]. Collectively, lower FA and white matter volumes in the callosum, cingulum and prefrontal areas could explain compromised inter-hemispheric communication in BD [9]. In BD, localized regions within anterior, middle, and inferior *fasciculi* showed lower FA, compared to their counterpart controls. Additionally, abnormalities in white matter microstructure were seen in segments of the occipital-parietal tracts, the vermis, brainstem, and spinal pathways. The signal detected within the uncinate fasciculus is of interest for BD pathophysiology.

This tract connects the orbitofrontal cortex and the anterior temporal lobes, and it plays a role in the amygdala-ventral prefrontal cortex system. While traditionally associated with the limbic system, the uncinate fasciculus may also be involved in episodic memory, language processing, and social-emotional processing [36]. A meta-analysis of TBSS studies reported lower FA in the uncinate fasciculus, as well as the internal capsule and corpus callosum [8]. The first meta-analysis of DTI studies in BD reported lower FA in the superior longitudinal (arcuate) and inferior *fasciculi*, inferior fronto-occipital gyrus, and posterior thalamic radiation, as well as the limbic right anterior and subgenual cingulate cortex [7]. Interestingly, significant group differences were found within posterior-cerebellar pathways in BD, with higher FA in localized regions in the BD group. These differences may represent a compensatory mechanism to the dysconnectivity observed across fronto-limbic and inter-hemispheric projections when accounting for the microstructural organization of the bundle. The cerebellum has been widely studied for its role in motor control, though through its extensive connections with association regions of the cortex the cerebellum is also integral to cognitive and affective processes. The presence of intact cell membranes and myelination may modulate anisotropy [37], so differences in FA may reflect neuroinflammation or changes in myelination. Other tensor-derived metrics such as medial (MD), radial (RD) and axial (AD) diffusivity, measuring other directional aspects of the diffusion profile, or multicompartment models such as NODDI, offer alternative metrics of white matter organization and future investigations should include these metrics in the context of BUAN tractometry to further characterize BD white matter abnormalities. We aimed to minimize differences in acquisition protocols throughout all quality checks, data processing, and statistical analyses, by checking signal-to-noise ratio, using a standardized pipeline, and treating site as random effect term. While we attempted to mitigate most known site effects in our analyses, it is important to note that we may have not fully controlled for the influence of all site-specific effects on tract reconstruction, which could potentially impact tract segmentation and the subsequent analyses. Using an advanced along-tract analytic method, BUAN, we conducted fine-scale spatial mapping of regional WM microstructure differences in BD, relative to controls, integrating data across sites in the largest study to date. The tracts implicated here connect regions with important functional roles in the regulation of emotions, motivation, decision-making, and cognitive control, which are impaired in BD. This work extends findings seen in a smaller sample [15], and highlights the complexity and functional specificity of fiber groups within complex large white matter bundles, and may guide future work characterizing WM in BD.

## Notes

### Competing Interest Statement

The authors have declared no competing interest.

### Summary of Updates

Revisions in the introduction and discussions. Results have not changed.

## References

[1] J. Rehm and K. D. Shield, “Global burden of disease and the impact of mental and addictive disorders,” Current Psychiatry Reports. Current Medicine Group LLC, vol. 1, 2019.

[2] S. O’Donoghue, L. Holleran, D. M. Cannon, and C. McDonald, “Anatomical dysconnectivity in bipolar disorder compared with schizophrenia: A selective review of structural network analyses using diffusion MRI,” J Affect Disord, vol. 209, pp. 217–228, 2017, doi: 10.1016/j.jad.2016.11.015.

[3] A. Mechelli, C. J. Price, K. J. Friston, and J. Ashburner, “Voxel-based morphometry of the human brain: methods and applications,” Curr Med Imaging, vol. 1, no. 2, pp. 105–113, 2005.

[4] G. Nortje, D. J. Stein, J. Radua, D. Mataix-Cols, and N. Horn, “Systematic review and voxel-based meta-analysis of diffusion tensor imaging studies in bipolar disorder,” J Affect Disord, vol. 150, no. 2, pp. 192–200, 2013, doi: 10.1016/j.jad.2013.05.034.

[5] P. Favre et al., “Widespread white matter microstructural abnormalities in bipolar disorder: evidence from mega- and meta-analyses across 3,033 individuals,” Neuropsychopharmacology, vol. 44, no. 13, pp. 2285–2293, 2019, doi: 10.1038/s41386-019-0485-6.

[6] L. Nabulsi et al., “Large-Scale Replication of Bipolar Disorder Dysconnectivity: A Diffusion MRI Analysis of 959 Individuals From the ENIGMA Bipolar Disorder Working Group,” Biol Psychiatry, vol. 89, no. 9, pp. S185–S186, 2021, doi: 10.1016/j.biopsych.2021.02.471.

[7] F. E. Vederine, M. Wessa, M. Leboyer, and J. Houenou, “A meta-analysis of whole-brain diffusion tensor imaging studies in bipolar disorder,” Prog Neuropsychopharmacol Biol Psychiatry, vol. 35, no. 8, pp. 1820–1826, 2011, doi: 10.1016/j.pnpbp.2011.05.009.

[8] L. Emsell and C. McDonald, “The structural neuroimaging of bipolar disorder,” International Review of Psychiatry, vol. 21, no. 4, pp. 297–313, 2009, doi: 10.1080/09540260902962081.

[9] L. Emsell et al., “White matter differences in euthymic bipolar I disorder: A combined magnetic resonance imaging and diffusion tensor imaging voxel-based study,” Bipolar Disord, vol. 15, no. 4, pp. 365–376, 2013, doi: 10.1111/bdi.12073.

[10] Y. Da, S. Mm, G. Sa, R. Ml, and P. Pj, “White matter abnormalities observed in bipolar disorder : a diffusion tensor imaging study,” vol. i, pp. 504–512, 2007.

[11] M. Wessa et al., “Microstructural white matter changes in euthymic bipolar patients: a whole-brain diffusion tensor imaging study,” Bipolar Disord, vol. 11, no. 8, pp. 504–514, 2009, doi: BDI718 [pii]\r10.1111/j.1399-5618.2009.00718.x.

[12] B. Q. Chandio et al., “Bundle analytics, a computational framework for investigating the shapes and profiles of brain pathways across populations,” Sci Rep, vol. 10, no. 1, p. 17149, 2020, doi: 10.1038/s41598-020-74054-4.

[13] B. Q. Chandio et al., “Microstructural changes in the white matter tracts of the brain due to mild cognitive impairment,” Alzheimer’s & Dementia, vol. 18, no. S5, Dec. 2022, doi: 10.1002/alz.065339.

[14] K. G. Schilling et al., “Aging and white matter microstructure and macrostructure: a longitudinal multi-site diffusion MRI study of 1218 participants,” Brain Struct Funct, vol. 227, no. 6, pp. 2111–2125, 2022.

[15] L. Nabulsi et al., “Along-Tract Statistical Mapping of Microstructural Abnormalities in Bipolar Disorder: A Pilot Study,” bioRxiv, p. 2023.03.07.531585, Jan. 2023, doi: 10.1101/2023.03.07.531585.

[16] B.C.M. Haarman et al., 2016. Diffusion tensor imaging in euthymic bipolar disorder – A tract-based spatial statistics study. J Affect Disord. 203:281–291.

[17] L. Nabulsi, L., et al., 2019. Bipolar Disorder and Gender are Associated with Fronto-limbic and Basal Ganglia Dysconnectivity: A Study of Topological Variance Using Network Analysis. Brain Connect.

[18] S. O’Donoghue S, L. Holleran L, et al., 2017. Anatomical dysconnectivity in bipolar disorder compared with schizophrenia: A selective review of structural network analyses using diffusion MRI. J Affect Disord. 209:217–228.

[19] G. Roberts, et al., 2018. Structural dysconnectivity of key cognitive and emotional hubs in young people at high genetic risk for bipolar disorder. Mol Psychiatry. 23:413–421.

[20] C. Vogelbacher, et al., 2018. The Marburg-Münster Affective Disorders Cohort Study (MACS): A quality assurance protocol for MR neuroimaging data. Neuroimage. 172:450–460

[21] A. Leemans, B. Jeurissen, J. Sijbers, and D. Jones, “ExploreDTI: a graphical toolbox for processing, analyzing, and visualizing diffusion MR data,” Proceedings 17^th^Scientific Meeting, International Society for Magnetic Resonance in Medicine, vol. 17, no. 2, p. 3537, 2009, [Online].

[22] J. Jeurissen et al., 2014. Multi-tissue constrained spherical deconvolution for improved analysis of multi-shell diffusion MRI data. NeuroImage, vol. 103, pp. 411–426, 2014, doi: 10.1016/j.neuroimage.2014.07.061.

[23] L.-C. Chang, D. K. Jones, and C. Pierpaoli, “RESTORE: Robust estimation of tensors by outlier rejection,” Magn Reson Med, vol. 53, no. 5, pp. 1088–1095, May 2005, doi: 10.1002/mrm.20426.

[24] E. Garyfallidis et al., “Dipy, a library for the analysis of diffusion MRI data, Frontiers in Neuroinformatics, vol. 8. 2014. [Online]. Available: https://www.frontiersin.org/articles/10.3389/fninf.2014.00008

[25] B. B. Avants, N. Tustison, and G. Song, “Advanced normalization tools (ANTS),” Insight J, vol. 2, no. 365, pp. 1–35, 2009.

[26] E. Garyfallidis, O. Ocegueda, D. Wassermann, and M. Descoteaux, “Robust and efficient linear registration of white-matter fascicles in the space of streamlines,” NeuroImage, vol. 117, pp. 124–140, 2015.

[27] F.-C. Yeh et al., “Population-averaged atlas of the macroscale human structural connectome and its network topology,” NeuroImage, vol. 178, pp. 57–68, 2018.

[28] E. Garyfallidis et al., “Recognition of white matter bundles using local and global streamline-based registration and clustering,” NeuroImage, vol. 170, pp. 283–295, 2018.

[29] J. D. Yeatman, R. F. Dougherty, N. J. Myall, B. A. Wandell, and H. M. Feldman, “Tract profiles of white matter properties: automating fiber-tract quantification,” PLoS One, vol. 7, no. 11, p. e49790, 2012.

[30] A. Yendiki et al., “Automated probabilistic reconstruction of white-matter pathways in health and disease using an atlas of the underlying anatomy,” Front Neuroinform, vol. 5, p. 23, 2011.

[31] B.Q. Chandio, “Advancing White Matter Tractometry of the Brain using Diffusion MRI and Machine Learning,” Indiana University, PhD Thesis, 2022.

[32] Y. Benjamini and Y. Hochberg, “Controlling the false discovery rate: a practical and powerful approach to multiple testing,” Journal of the Royal Statistical Society: Series B (Methodological), vol. 57, no. 1, pp. 289–300, 1995.

[33] L. Nabulsi et al., “Normalization of impaired emotion inhibition in bipolar disorder mediated by cholinergic neurotransmission in the cingulate cortex,” Neuropsychopharmacology, vol. 47, no. 9, pp. 1643–1651, 2022.

[34] C. R. K. Ching et al., “What we learn about bipolar disorder from large-scale neuroimaging: Findings and future directions from the ENIGMA Bipolar Disorder Working Group,” Hum Brain Mapp, no. May, pp. 1–27, 2020, doi: 10.1002/hbm.25098.

[35] A. Perry, G. Roberts, P. B. Mitchell, and M. Breakspear, “Connectomics of bipolar disorder: a critical review, and evidence for dynamic instabilities within interoceptive networks,” Mol Psychiatry, 2018, doi: 10.1038/s41380-018-0267-2.

[36] R. J. von der Heide, L. M. Skipper, E. Klobusicky, and I. R. Olson, “Dissecting the uncinate fasciculus: disorders, controversies and a hypothesis,” Brain, vol. 136, no. 6, pp. 1692–1707, 2013.

[37] Phillips JR, Hewedi D.H., Eissa A.M., Moustafa A.A., “The cerebellum and psychiatric disorders,” Front Public Health, vol. 3, p. 66, 2015.

